# 3D pathology-guided microdissection

**DOI:** 10.1101/2025.11.20.689586

**Authors:** Huai-Ching Hsieh, Gan Gao, Qinghua Han, David Brenes, Elena Baraznenok, Renao Yan, Robert Serafin, Kevin W. Bishop, Rui Wang, Eric Q. Konnick, Colin C. Pritchard, Sandy Figiel, Freddie C. Hamdy, Ian G. Mills, Nicholas P. Reder, Deepti M. Reddi, Thomas G. Paulson, William M. Grady, Jacob E. Valk, Lawrence D. True, Michael C. Haffner, Srinivasa R. Rao, Dan J. Woodcock, Jonathan T.C. Liu

## Abstract

Traditional micro- and macro-dissection techniques enable the extraction of localized regions in thin tissue sections for molecular analysis. Despite the growing use of 3D microscopy, analogous methods for volumetric microdissection are lacking. We have developed a 3D microdissection method based on computer numerical controlled (CNC) milling integrated with open-top light-sheet microscopy. We demonstrate the ability to study tumor evolution along convoluted 3D branching architectures, which is inaccessible to 2D methods.

---

Micro- and macro-dissection techniques enable the isolation of localized tissue regions under image guidance. The extracted material is subsequently used for various omics assays, enabling the integrated analysis of morphological and molecular data^1, 2^. These methods have been applied to a wide range of tissue types and investigations, including the clonal evolution of cancers^3, 4^ and the study of region-specific properties in heterogeneous tissue microenvironments^5^. A simple method commonly performed in clinical settings is manual macrodissection, which involves scraping tissue from slides (mm–cm scale)^6^. Laser capture microdissection (LCM), used for basic investigations, relies on precise laser control to extract defined tissue regions of interest (μm–mm scale)^1,7^. Despite their complementary strengths, both approaches are fundamentally limited to thin tissue sections and are guided by 2D microscopy.

In recent years, spatial omics techniques have transformed our ability to link molecular and spatial information^8–14^. However, they are largely confined to thin 2D tissues sections and are typically incompatible with comprehensive molecular-profiling, such as high-depth whole-genome sequencing^8^. There is a growing appreciation for the importance of 3D information for understanding biological processes^15–17^, such as tumor–stroma interfaces and invasive growth patterns (Fig. 1a). Emerging 3D spatial omics methods provide volumetric context but lack broad molecular coverage and are limited in tissue depth (<300 um)^12, 13^. There remains an unmet need for technologies to enable integrated spatial-molecular analyses with comprehensive high-depth omics at the cell-population level in conjunction with high-resolution 3D morphological information across mm-cm length scales.

**Figure 1.**
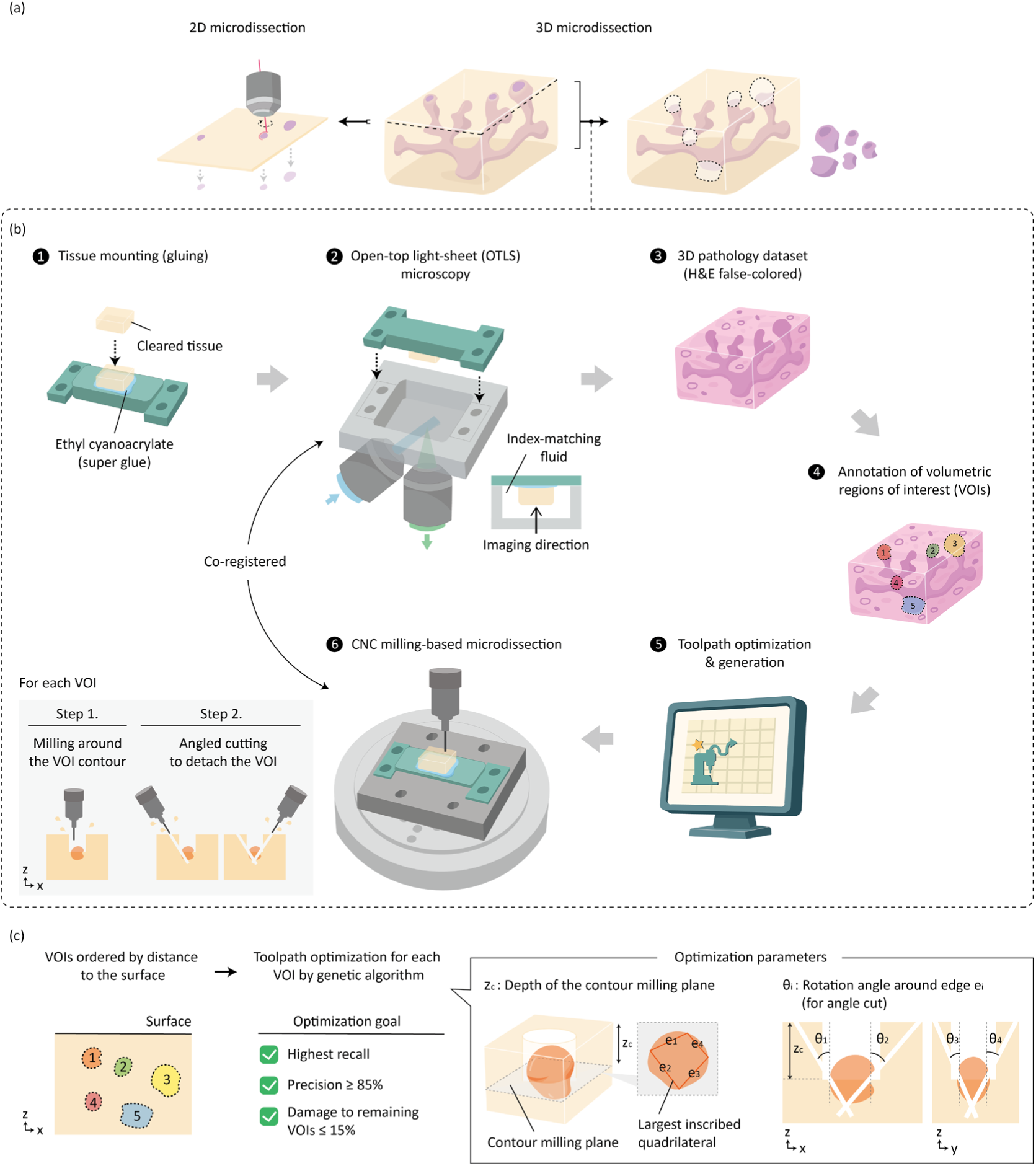
(a) Traditional 2D microdissection isolates regions from thin sections and cannot capture 3D morphology (e.g., branching tumors), whereas 3D microdissection enables comprehensive characterization and extraction from 3D structures. (b) Workflow of 3D pathology-guided microdissection (3DPM). (c) Toolpath optimization strategies and parameters for maximizing recall and precision while minimizing damage to other VOIs.

We present a 3D microdissection method that first acquires volumetric tissue images using open-top light-sheet (OTLS) microscopy^18, 19^, followed by computer numerical control (CNC) milling–based extraction of volumetric regions of interest (VOIs) based on 3D annotations. Accurate localization of VOIs is enabled via a co-registration process between the 3D microscopy and milling systems. A key challenge in 3D microdissection is achieving spatially precise detachment of VOIs in all three dimensions, particularly for closely spaced VOIs. We address this challenge by designing a workflow to generate oblique CNC milling paths that are computationally optimized to maximize recall (which reflects tissue yield) and precision (which reflects sample purity) for a collection of desired/annotated VOIs. In practice, the current system can achieve dissection of volumes down to ~1 mm laterally and ~300 µm axially.

A major application of traditional 2D LCM on thin histology sections has been to study tumor phylogeny, in which subclonal relationships are reconstructed amongst distinct tumor regions^3, 4, 20^. However, due to the lack of information on 3D spatial connectivity and tumor architecture, important questions remain unresolved, such as how tumors evolve and disseminate along spatially complex branched trajectories, and how these spatial branches relate to clonal diversification. While earlier 3D reconstruction efforts based on 2D data have provided valuable insights^21, 22^, comprehensive spatial phylogenetic mapping was still lacking due to technological limitations. Here, we demonstrate that 3D pathology-guided microdissection (3DPM) has the potential to address these limitations and to enable new insights into cancer evolution through “3D spatial tumor phylogeny.”

The 3DPM workflow (Fig. 1b) begins with mounting optically cleared tissue onto a custom holder to ensure positional stability during both imaging and microdissection. The holder fits into matching slots in the OTLS-microscopy and CNC systems that enable repeatable placement on both devices (Extended Data Fig. 1). Co-registration, i.e. computing the coordinate transformation between systems, is achieved in advance by milling a reference pattern onto the holder and imaging it (Extended Data Fig. 2a-c, e). After OTLS microscopy imaging of a tissue sample, a 3D pathology dataset is generated, where established false-coloring algorithms may be applied to mimic the appearance of H&E histology to facilitate interpretation by end users^23, 24^. Based on manually annotated VOIs, the CNC toolpaths are optimized and generated. As a subtractive manufacturing process, traditional CNC milling is designed to engrave or remove material rather than to preserve small, spatially defined structures. Here, we introduce a novel toolpath strategy tailored for microdissection: first, the tissue is removed above a plane that encloses the VOI; second, angled cuts are applied along four quadrants to fully detach the VOI from the base. These microdissection parameters—the depth of the first cutting plane and the four cutting angles (with respect to the vertical axis)—are optimized to maximize recall for each VOI while minimizing collateral damage to neighboring VOIs that have yet to be extracted (Fig. 1c, Extended Data Fig. 3, 4).

To validate the 3D microdissection workflow, we applied it to multiple tissue types, including cartilage, head and neck, prostate, and breast specimens, which collectively exhibit a wide range of Young’s moduli (stiffness) after optical clearing (Extended Data Fig. 2d). For each tissue type, a specimen with a size of roughly 1 cm laterally and 1.3 mm axially was imaged, and six VOIs of varying size, shape, and spatial distribution were annotated and microdissected (average volume: 0.72 mm³; Extended Data Fig. 2f). To quantify recall and precision metrics, extracted tissues were re-imaged and compared with the original annotations. Figure 2a shows representative comparisons between microdissected samples and their original annotations, showing good qualitative correspondence. Quantitative evaluation further demonstrates robust recall and precision across tissue types, with average recall and precision metrics of 65.1% and 63.8%, respectively. Even in low-stiffness tissue, such as normal breast, VOIs could be successfully microdissected (Extended Data Fig. 2g).

**Figure 2.**
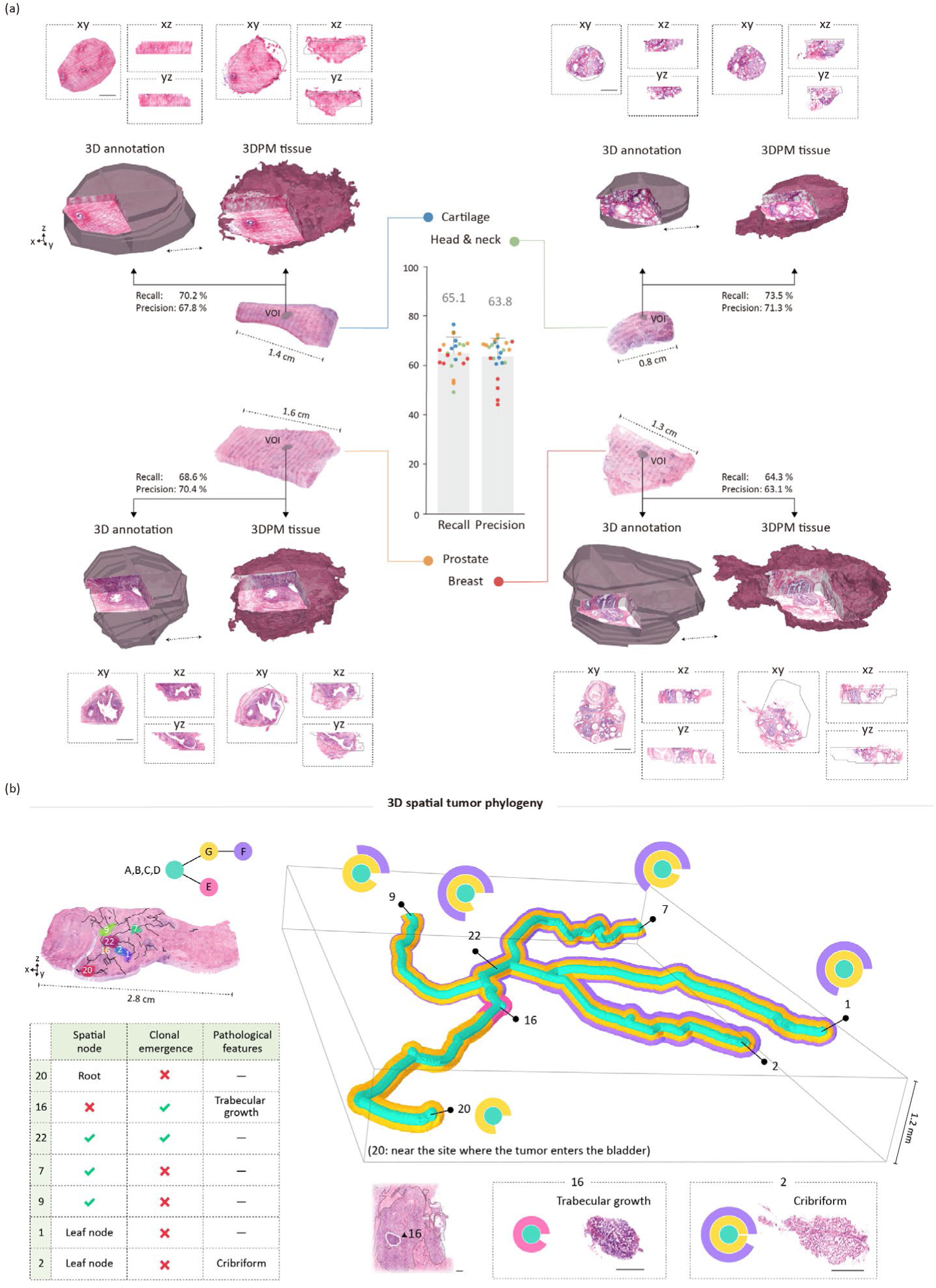
(a) Quantification of 3DPM performance and comparisons between VOI annotations and microdissected tissues for four tissue types. The gray contours in the cross-sectional views indicate the original VOI annotations. Cut-away views are provided at the bottom left quadrant of each of the VOIs (pre- and post-dissection imaging), with corresponding 2D cross-sectional views shown at various cut planes; scale bar (with and without black arrows) = 500 μm). (b) Upper left: reconstructed phylogenetic tree and 3D rendering of the tissue block overlaid with the skeletonized gland network. Seven VOIs are shown along this branched 3D network. Lower left: table summarizing spatial nodes, phylogenetic points of interest, and pathological characteristics for each VOI. Right: reconstructed 3D spatial tumor phylogeny tree and corresponding sunburst plots, with the center corresponding to the ancestral clone and each outward ring representing a subsequent clone in the phylogenetic tree^4^. The thickness (3D tree) or the angular span (2D plot) of each colored band represents the cancer cell fraction for that clone (360°: 1). (scale bar = 500 μm)

We next applied 3DPM to study a prostate cancer case involving the bladder with a highly branched architecture. Tumor segmentation was manually performed under the guidance of a board-certified anatomic pathologist subspecialized in GU Pathology. A 3D skeletonization approach was used to extract the main structural trajectory of the tumor. To investigate the clonal relationships between distinct spatial branches within the tumor, seven VOIs were selected for microdissection and whole-genome sequencing. The phylogenetic tree was reconstructed from somatic SNVs, adjusting for local copy-number states. These sites encompassed the skeleton root near the site where the tumor enters the bladder, major branching nodes, and terminal leaf nodes forming multiple branches that extend in divergent 3D directions. Additionally, a region was demarcated with distinct pathological features (VOI 16; Fig. 2b) located near a highly branching node.

We established the 3D spatial tumor phylogenetic tree by inferring possible dissemination routes of subclones between VOIs based on their cancer cell fractions (CCFs, i.e., the proportion of cancer cells within a sample that harbors a given set of mutations), and overlaid these routes on the 3D tumor skeleton (Fig. 2b, Supplementary Video 1). The branch lengths reflect physical distances between regions in the tissue. With only 2D spatial information of VOIs, dissemination paths cannot be uniquely inferred, as one phylogenetic relationship may correspond to multiple spatial paths with different biological implications (Extended Data Fig. 6). In contrast, 3D spatial phylogeny analysis reveals a most likely and spatially coherent dissemination path. In addition, comparison of the spatial branching and phylogenetic trajectories reveals diverse relationships: in this case, a single site acts as a node in both trees, whereas other nodes show divergence only in 3D spatial trajectories or phylogeny. These results indicate that spatial branching and clonal emergence are not always coupled, highlighting the spatial and evolutionary complexity of tumors and the necessity of integrating spatial and phylogenetic analyses for a full understanding of tumor evolution.

Despite its capabilities, 3DPM has several current limitations. In order to microdissect VOIs in deeper parts of the tissue, destructive removal of some tissue above and around the VOI is unavoidable — but our algorithm minimizes such collateral damage. Another limitation relates to sample preprocessing: dehydration-based clearing enhances tissue stiffness^25, 26^, which improves the precision of CNC milling, but is dependent upon tissue type and is not compatible with some stains and antibodies. Future work will explore alternative strategies to increase tissue stiffness while maintaining optical transparency and compatibility with various staining options. In addition, the collection of microdissected tissue is currently performed manually. Integration of an automated collection system is planned to enhance throughput and ease of use. Finally, while the current system can achieve a minimum lateral microdissection size of ~1 mm, further miniaturization with finer milling tools (and/or stiffer tissue) could enable the extraction of smaller VOIs.

Several prior approaches have been attempted to integrate a degree of volumetric and molecular information. One category relies on serial sectioning and 3D reconstruction, such as performing 2D spatial omics at 100-μm depth intervals^21^, or applying LCM to serial sections followed by 3D reconstruction^22^. These methods provide high-resolution molecular data and partial 3D context, but with intrinsic limitations. Physical sectioning can distort or destroy delicate structures—such as invasive fronts near nerves or interconnected glands/crypts. When performed serially for 3D reconstruction, such processes are labor-intensive and can introduce reconstruction artifacts (e.g., cracks and deformations) as well as co-registration errors between sections, making accurate 3D spatial relationships difficult to establish. Such artifacts may be especially confounding when attempting to confirm the connectedness (or lack of connectivity) between convoluted/branching microstructures. Moreover, for spatial omics techniques that rely on sparse sampling, molecular sampling is typically based on 2D morphological representations that can misrepresent 3D structures ^18, 27, 28^. In contrast, 3DPM is guided by a full 3D morphological context provided by high-throughput non-destructive 3D microscopy, allowing for the extraction of intact tissue volumes for molecular profiling (Extended Data Fig. 6).

Co-registering volumetric imaging with a mechanical device has been demonstrated in previous works. One study integrated a robotic vacuum-aspiration system with 3D microscopy to extract cell populations^29^. However, since suction was applied at various points without delineation of a volumetric boundary for the region of interest, the size and shape of the extracted tissue/cells could not be precisely controlled. In addition, suction-based extraction resulted in tissue that was deformed or fragmented.

Here, we have demonstrated a novel CNC milling–based microdissection approach to enable volumetric cutting with structural preservation. By introducing an innovative pipeline for 3D tissue extraction, our method addresses several critical needs. First, 3DPM employs volumetric annotation to optimize the sampling fraction, ensuring that molecular profiles faithfully represent the defined region of interest. Preserving the 3D structure/morphology of the extracted tissues further allows quantitative performance evaluation, providing a rigorous means to validate and optimize microdissection performance (e.g. recall and precision). This also enables potential integration with high-plex spatial omics techniques, which can be applied to an array of rare or small VOIs microdissected from a large sample. Collectively, these capabilities position 3DPM as a novel framework that unites 3D pathology (and related 3D imaging techniques) with comprehensive omics, providing new insights that are difficult to achieve with existing spatial-biology tools.

## Methods

### Tissue preprocessing and mounting

Tissue preprocessing followed a previously established 3D pathology protocol^18, 24^, with modifications according to tissue type. In this study, prostate tissues used for validation were stored in ethanol; breast, head and neck, and chicken cartilage tissues were obtained fresh; and bladder tissue with prostate cancer was obtained as a FFPE block. Formalin-fixed paraffin-embedded (FFPE) tissues were first heated at 70 °C for 1 hour to melt the paraffin, then incubated in Histo-Clear II (HS-202, National Diagnostics) at room temperature for 48 hours to achieve complete deparaffinization. Residual solvent was removed by a 100% ethanol wash. Fresh tissues were fixed overnight in formalin (3800602, Leica Biosystems) and rinsed with 70% ethanol. Samples stored in 70% ethanol proceeded directly to staining and clearing. All samples were stained using TO-PRO-3 (10710194, Thermo Fisher Scientific) and Eosin Y (3801615, Leica Biosystems) – fluorescent analog of H&E – in 70% ethanol adjusted to pH 4 with 10 mM NaCl for 48 hours. Samples were then dehydrated in 100% ethanol and incubated overnight in ethyl cinnamate (Eci; A12906, Thermo Fisher Scientific) for optical clearing. After clearing, samples were mounted onto a custom Hi-Vex tissue holder using ethyl cyanoacrylate superglue (AD112, 3M Science) and allowed to stabilize for 30–60 minutes before imaging.

### Open-top light-sheet (OTLS) microscopy, image processing, and 3D annotation

All samples were imaged using a custom-built axially swept OTLS microscope^19^ (**Fig. 1b** and **Extended Data Fig. 1**). The sample holder was inverted and positioned within a matching slot on a custom Hi-Vex imaging chamber, with ECi used as the index-matching medium (refractive index = 1.56). TO-PRO-3 fluorescence was excited at 638 nm and collected using a long-pass emission filter (BLP01-647R-25, Semrock), while Eosin fluorescence was excited at 561 nm and collected using a band-pass emission filter (FF01-618/50-25, Semrock). Volumetric images were downsampled to 3.9 µm/voxel for the 3D microdissection validation dataset and to 2.9 µm/voxel for the bladder tissue used in phylogenetic analysis. Image tile stitching and fusion were performed using the BigStitcher plug-in in Fiji^24, 30^. A previously established H&E false-coloring algorithm, based on the Beer–Lambert law of absorption, was applied to the two-channel fluorescence images to simulate the appearance of standard H&E images^23^. 3D annotations were generated by manually labeling image sections at 50-µm intervals on the false-colored 2D image stack using QuPath^31^, followed by 3D upsampling to generate continuous annotated volumes. 3D skeletonization of the tumor mask was performed using the *Kimimaro* Python package (version 5.0.0).

### Co-registration between OTLS and 5-axis CNC milling-based microdissection system

A desktop 5-axis CNC milling machine (Pocket NC V2-50CHK, Penta Machine) was used for 3D microdissection^32^, with a custom slot-matching adapter to ensure repeatable placement of the tissue holder. Prior to co-registration, the “zero working coordinate” of the CNC was manually set to one corner of the tissue holder. Co-registration was performed by engraving a target pattern on the tissue holder using a 500-µm diameter endmill (PD30A, SainSmart); the toolpath was generated in Autodesk Fusion 360 software (Autodesk Inc.). After engraving, the holder was imaged in ECi, with fluorescein dye added, to visualize the engraved pattern (**Extended Data Fig. 2a**). The acquired OTLS image was then aligned to the G-code toolpath, allowing for the derivation of a formula to transform the OTLS stage coordinates to the CNC working coordinates. To quantify co-registration errors, we engraved six cross-shaped patterns on the tissue holder at randomly selected OTLS stage coordinates. The average co-registration errors were 31.2 µm, 28.8 µm, and 26.9 µm in the X, Y, and Z directions, respectively (**Extended Data Fig. 2b**). We further assessed co-registration performance on optically cleared tissue, representing real-world conditions. Four tissue types with varying stiffness—prostate, head and neck, breast, and cartilage—were tested. For each tissue, multiple cross-shaped patterns (1 mm in width and height) were engraved at various depths using a 200-µm diameter endmill (PD30A, SainSmart) and then imaged with OTLS microscopy. Tissue-level co-registration errors were then quantified by comparing the imaged/measured center positions and milling depths of the engraved crosses with the theoretical (intended) values. The average errors were measured as 119 µm, 56.8 µm, and 108.9 µm in the X, Y, and Z directions, respectively, demonstrating robust performance across tissue types (**Extended Data Fig. 2c**).

The lower error observed in the Y direction may be attributed to the geometry of the tissue holder, as the Y axis corresponds to its longer edge, which is potentially less susceptible to variabilities in manufacturing and enables more-precise/reproducible placement of the tissue holder within its receptacle along that edge. As expected, greater errors were observed in softer tissues such as breast due to tissue deformation (**Extended Data Fig. 2e**). In particular, while XY-plane errors were relatively consistent across tissue types, Z-direction errors were higher in softer tissues, potentially due to compression-induced vertical deformations (i.e., the actual milling depth was shallower than the designed depth). We therefore recommend experimentally determining tissue-specific Z-offsets prior to 3D microdissection and applying appropriate depth corrections. The derived transformation formula was used for subsequent 3D microdissection experiments.

### 3D microdissection toolpath optimization

After 3D annotation of VOIs, their microdissection order is first determined based on the distance to the tissue surface, with VOIs closer to the surface being prioritized. For each VOI, toolpath optimization is performed using a hybrid strategy combining geometry-informed random search with a genetic algorithm (GA)^33^ (**Fig. 1; Extended Data Fig. 3**). Each candidate solution consists of five parameters: *z*_*c*_, which defines the depth of the contour milling plane, and four angles 𝜃_*i*_ (*i* = 1 − 4), which describe the tilt of the cutting planes. These cutting planes extend from the four edges of a “maximum inscribed quadrilateral” fitted to the 2D VOI cross-section at plane *z*_*c*_. For concave cross-sections, this quadrilateral is not required to be strictly inscribed.

The GA is initialized with 80 individuals (i.e., parameter sets). The contour milling-plane depth *z*_*c*_ is randomly selected near the vertical midpoint *z*_*median*_ of the VOI, within the VOI’s vertical bounds:

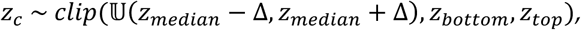

where Δ = 387 μm, *clip* (*x*, *a*, *b*) denotes constraining *x* to the interval [*a*, *b*], and 𝕌(*a*, *b*) represents a continuous uniform probability distribution over the interval [*a*, *b*].

Each angle 𝜃_*i*_ is then randomly initialized within ±30° of a heuristic angle 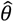, with a constraint to avoid extreme cuts:

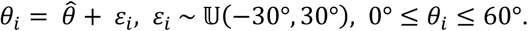

The heuristic angle 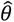 is computed as:

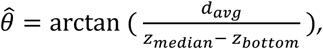

where *d*_*avg*_ is the average lateral distance from the centroid of the fitted quadrilateral to its four edges (at plane *z*_*median*_).

For each individual, a simulated 3D microdissection is performed: in the first step, contour milling is simulated by removing all off-target material within a vertical cylinder centered on the 2D cross section of the VOI at plane *z*_*c*_, using the minimum enclosing circle plus a 0.4 mm radial buffer (total radius *r*_*t*_); in the second step, four angled cuts are applied. Each cutting edge is extended beyond its original lateral endpoints by at least 50% of its length on both sides, providing a buffer to ensure full detachment of the VOI from its base.

Candidates that fail to completely detach the VOI are assigned zero recall and precision. Successful candidates are further evaluated to assess the maximum damage they create to the remaining, unextracted VOIs. A fitness function is defined as the recall of the extracted VOI on the condition that it meets a set of criteria for both precision and damage:

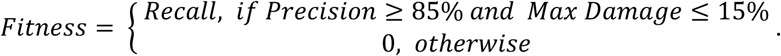

Recall and precision are computed based on the volumetric overlap between the target VOI and the successfully detached tissue volume:

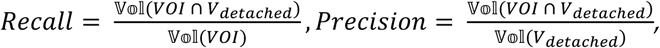

where 𝕍𝕠𝕝(·) denotes a 3D volume, and *V*_*detached*_ refers to the subset of the simulated microdissection region that is successfully separated from the base.

Maximum collateral damage to the remaining VOIs is defined as:

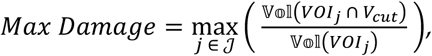

where *V*_*cut*_ includes all of the material removed by the simulated milling and angled cuts, and 𝒥 denotes the set of VOIs that have not yet undergone simulated dissection. We further evaluate whether *V*_*cut*_ creates multiple disconnected components within each *VOI*_*j*_. If so, we conservatively assume that only the largest component is preserved, and that all remaining fragments may be treated as damage.

After evaluating the initial population (generation 0), the best-performing individual is selected. If its recall exceeds 30%, it is considered an acceptable final solution and no further iterations are performed (to reduce computational costs). Otherwise, the GA is allowed to evolve for up to five additional generations (six total). In practice, acceptable solutions are achieved for most VOIs within the initial random pool.

### 3D microdissection toolpath generation

The optimized parameters for each VOI are subsequently used to generate the G-code that controls the 5-axis CNC system. The semi-automated workflow consists of two main steps. First, custom Python scripts are used to generate four computer-aided design (CAD) models representing geometries derived from the optimization output (**Extended Data Fig. 4a, b**).

1. *CAD*_1_: A full bounding box model extending from the base to the tissue surface.
2. *CAD*_2_: *CAD*_1_ with a vertical cylindrical cut (radius = *r*_*t*_) down to the top plane of the VOI.
3. *CAD*_3_: *CAD*_1_ with a vertical cylindrical cut (radius = *r*_*t*_) down to the plane *z*_*c*_, plus the portion of the VOI above plane *z*_*c*_
4. *CAD*_4_: A set of four bodies, each defined by subtracting a tilted cutting-plane box from *CAD*_1_ at an optimized angle 𝜃_*i*_. The thickness of each tilted box matches the endmill diameter. In this study, a 200-µm endmill was used for all 3D microdissections (PD30A, SainSmart).

These models are then imported into the computer-aided manufacturing (CAM) module of Autodesk Fusion 360 to define toolpaths. The work coordinate system (WCS) origin is set to the corner of the tissue holder that is used to define the zero coordinate during co-registration. Machining parameters—including spindle speed, cutting feed rate, and plunge feed rate—were experimentally optimized to minimize tissue deformation with the final values being 24,000 rpm, 50 mm/min, and 30 mm/min, respectively. The toolpath generation proceeds as follows (*CAD*_1_ is used to define the stock model) (**Extended Data Fig. 4c, Supplementary Video 2**):

1. *CAD*_2_ defines a 2D Pocket path to remove material above the VOI.
2. *CAD*_3_ is first used to generate a 3D Flow operation to engrave the portion of the VOI above plane *z*_*c*_.
3. Several 2D Contour paths are then applied to the cross-section at plane *z*_*c*_, with radial stock-to-leave offsets ranging from 0 mm to 0.3 mm.
4. *CAD*_4_ defines the angled cuts for VOI detachment. Here, 2D Slot operations are applied. To minimize tissue deformation, the four angled cuts are performed sequentially and cyclically, where each cut removes a small portion of the tissue. The first round of cuts reaches near the *z*_*c*_ plane. Subsequent paths incrementally deepen the cuts by 0.1 mm one edge at a time (cyclically) until the bottom of all four tilted planes are reached. A custom python code (see Code Availability) is used to streamline this process.

Final toolpaths are post-processed and exported as machine-specific G-code. This two-step workflow—CAD generation followed by CAM programming—is adopted instead of directly generating G-code for two key reasons. First, toolpath generation for 5-axis CNC milling is complex and typically requires professional CAM software to handle variable tool orientation, cutting logic, collision avoidance, and safety constraints. Second, G-code is a low-level machine language that depends heavily on the specific hardware and controller, making it difficult to generalize or maintain across systems.

### 3D microdissection operation

Once optimized, the G-code files are executed on the 5-axis CNC system to perform microdissection. At the end of each operation (four steps in **Extended Fig. 4c**), the endmill is cleaned with a brush to remove residual tissue debris, ensuring precise cutting. Microdissected tissues are manually collected upon visual confirmation of successful detachment of the VOIs from their tissue base. To prevent cross-VOI contamination, the endmill is rinsed with 100% ethanol between VOIs. For different cases (i.e., different tissue blocks), the endmill is replaced entirely to avoid cross-case contamination.

### Validation and quantification of 3D microdissection performance

For validation, the isolated tissues were re-imaged using OTLS after 3D microdissection. Manual rotation and transformation were applied to align the original annotation with the resulting microdissected tissue. Tissue contours were segmented from Eosin Y channel images (cytoplasmic signal) using Otsu thresholding followed by manual correction. Two metrics were calculated: recall, defined as 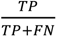, and precision, defined as 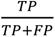. Here, TP (true positive) denotes the region that was both annotated and successfully microdissected; FN (false negative) refers to the annotated region that was missed; FP (false positive) represents regions that were not originally annotated but were nonetheless microdissected. Since the definition of TN (true negative) is arbitrary in this context, accuracy was not reported. As the quantification of 3D microdissection performance requires morphological mapping of microdissected tissues back to originally annotated regions, only VOIs with recognizable structures were selected during the initial annotation process. These VOIs spanned different depths within the tissue block to enable an assessment of average performance. Note that some structural deformation may occur during the microdissection process.

### DNA extraction of 3D microdissected samples

For samples used in phylogenetic analyses, 3D microdissected tissues were washed sequentially in 100% and 70% ethanol, each for one day. The tissues were then rehydrated and washed in phosphate-buffered saline (PBS). DNA extraction was performed by the Biospecimen Processing & Biorepository Core at Fred Hutchinson Cancer Center using a modified protocol based on the QIAamp DNA FFPE Tissue Kit (Qiagen). Briefly, tissues were first digested with Buffer ATL and Proteinase K at 56 °C overnight; a tissue grinder pestle was optionally used to enhance tissue lysis. Subsequent steps followed the manufacturer’s standard protocol. DNA concentration and quality were assessed using Qubit (Thermo Fisher Scientific) and TapeStation (Agilent Technologies), respectively.

A FFPE prostate tissue block was used to validate DNA quality and integrity following the 3DPM workflow and the modified DNA extraction protocol. Four 3-mm punches were randomly obtained: two were processed as standard FFPE curls with DNA extracted using the QIAamp DNA FFPE Tissue Kit, while the other two underwent the complete workflow for 3D pathology tissue preprocessing and imaging, followed by the modified DNA extraction protocol. DNA integrity was assessed using the DNA Integrity Number (DIN), and PCR performance was evaluated using *GAPDH* and *COL2A1* primers (target lengths: 228 bp and 274 bp; Hs00828901_CE and Hs00570192_CE, Thermo Fisher Scientific). Comparable DIN values and PCR amplification performance were observed between the two groups (**Extended Data Fig. 5**).

### Library preparation and whole-genome sequencing (WGS)

After genomic DNA extraction, library preparation and whole-genome sequencing were performed by the Genomics & Bioinformatics core at the Fred Hutchinson Cancer Center. WGS libraries were prepared using the IDT xGen cfDNA & FFPE Library Prep Kit (Integrated DNA Technologies), following the manufacturer’s instructions. Library quality was assessed using a TapeStation system (Agilent Technologies), and sequencing was performed on 25B-lane NovaSeq X Plus using 150-bp paired-end reads. The mean coverage across 7 tumor samples was 40X, and 18X for the matched normal sample. In this study, the adjacent muscle region was used as the normal sample.

### WGS data preprocessing

Raw FASTQ data were first trimmed to remove adapter sequences using BBDuk^34^ (version 39.24), and then aligned to the hg38 reference genome using BWA-MEM^35^ (version 0.7.19-r1273). Aligned reads were coordinate-sorted using Samtools^36^ (version 1.21), and duplicate marking was performed with MarkDuplicates from the Genome Analysis Toolkit (GATK, version 4.6.2.0)^37^. Read quality was assessed before and after trimming using FastQC^38^ (version 0.12.1). The preprocessing pipeline was adapted from Rao et al. (2024), with modifications for reference genome version and library-specific adjustments^4^.

### Single nucleotide variant (SNV) calling

BAM files with duplicate marking were further processed using Base Quality Score Recalibration (BQSR) to recalibrate base quality scores. Somatic variant calling was then performed using GATK Mutect2 in multi-tumor mode, specifically for SNV detection. The resulting variant calls were filtered using GATK FilterMutectCalls with an appropriate F-score parameter to balance sensitivity and specificity and to remove low-frequency or low-confidence SNVs prior to subclonal analysis. Manual filtering based on variant frequency, coverage, and mutant coverage was also applied on top of FilterMutectCalls to further remove low-confidence SNVs. The SNV calling pipeline was adapted from Rao et al. (2024), with modifications for reference genome version and library-specific adjustments^4^.

### Copy number alterations (CNA) calling

CNAs were called using the Battenberg R package^39^ (version 2.9.9.9000) on BAM files processed with base quality score recalibration (BQSR). To assess the quality of CNA calling, we adapted the quality check (QC) method described in Cornish et al. (2024)^40^. Briefly, DPClust (version 2.2.8) was first run in single-sample mode, using SNVs and Battenberg-derived CNAs. Each sample was evaluated for the presence of a clonal cluster, defined as the cluster with the largest number of SNVs and a cancer cell fraction (CCF) between 0.95 and 1.05. Samples meeting this criterion were designated as DPClust-pass. In parallel, CNAqc (version 1.1.2) was applied using the Battenberg-estimated purity. Samples were marked as CNAqc-pass if they passed the built-in QC and the purity difference was less than 5%. If either check failed, Battenberg CNA calling was repeated using revised estimates of purity and ploidy. The CNA calling pipeline was adapted from Rao et al. (2024), with modifications to the reference genome version, library-specific parameters, and QC procedures^4^.

### Phylogenetic analysis

DPClust-3p^41^ (version 1.0.8) was first applied in each sample to generate cancer cell fraction (CCF) estimates based on SNVs and CNAs. Multi-sample clustering was then achieved using the BayesianGaussianMixture function in Scikit-learn Python package^42^ (version 1.7.0) with 1000 iterations, after which clusters were refined by visual inspection, with merging or splitting applied as appropriate. For each resulting subclone, the clonal fraction was defined as the median CCF of all SNVs assigned to that cluster. The clustering pipeline was adapted from Rao et al. (2024)^4^, with modifications to the reference genome version and library-specific parameters. Phylogenetic trees were reconstructed following the sum rule and crossing rule^20^. The sum rule mentions that the sum of the CCFs of the daughter clones must be less than that of the parent clone. The crossing rule states that in multi-region sequencing (i.e., multiple VOIs in this study), if clones B and C are both daughter clones of A, and there exists one sample in which CCF(B) > CCF(C) and another sample in which CCF(C) > CCF(B), then B and C must be placed on different branches. When contradictions arose between these rules, the solution with the fewest violations was selected.

### Laser-capture microdissection (LCM)

Due to the increased complexity of cutting along 3D contours compared to 2D, LCM was performed on a tissue section cut from the top of the FFPE block used for the 3D phylogenetic analysis of bladder-involved prostate cancer. This allowed for a comparison of tumor purity between regions microdissected using 2D and 3D approaches. The LCM slide was prepared by sectioning a 10-µm slice and mounting it onto a polyethylene naphthalate (PEN) membrane slide (LCM0521, Thermo Fisher Scientific). Slide preparation was carried out by the Experimental Histopathology Core at the Fred Hutchinson Cancer Center. The LCM experiment was then performed using a Leica LMD6 laser microdissection system under the guidance of an LCM specialist. H&E staining was performed prior to LCM to visualize histological structures. Five tumor regions were selected based on the projection of 3D VOIs from the 3D microdissection onto the 2D slide (**Extended Data Fig. 7a**), and one adjacent normal region was also dissected. Cell lysis and DNA purification were adapted from Ellis et al. (2021)^43^ and carried out by the Biospecimen Processing & Biorepository Core at the Fred Hutchinson Cancer Center. Briefly, collected tissue fragments were lysed using the Arcturus PicoPure DNA Extraction Kit (KIT0103, Thermo Fisher Scientific), following the manufacturer’s instructions. DNA was then purified using AMPure XP magnetic beads (Beckman Coulter Life Sciences). After bead removal, DNA concentration and fragment size were assessed using Qubit and TapeStation, respectively. Library preparation and whole-genome sequencing (WGS) were performed as described in the previous sections. The mean coverage across the five tumor LCM samples was 31X, and 23X for the matched normal sample. WGS preprocessing, BQSR recalibration, and the Battenberg pipeline were subsequently applied to estimate tumor purity.

**Extended Data Fig. 7b** compares tumor purity between the 2D LCM and 3DPM methods. While the 3DPM method shows a slight decrease in purity, the difference between the two methods is not statistically significant based on either the two-tailed independent t-test (p = 0.54) or the two-tailed paired t-test (p = 0.28, using regions 16_22’ and 1_2’ in 2D LCM paired with VOIs 16, 22 and VOIs 1, 2, respectively), indicating comparable performance. Two lower-purity VOIs observed in 3DPM may have resulted from complex tumor morphologies and/or indistinct histological features. To provide a control for tumor evolutionary inference, a joint analysis of 2D LCM and 3DPM samples was performed, including joint SNV calling and clustering, as shown in **Extended Data Fig. 7c**. This comparison between methods demonstrated good agreement between the two approaches, while also revealing better clustering results in 3DPM — likely because it enables the collection of larger and more representative tissue volumes compared to the limited material obtainable with 2D LCM. (**Extended Data Fig. 7d,e**).

### Young’s modulus measurement of optically cleared tissue

After optical clearing, tissues were trimmed into a rectangular shape. The Young’s modulus was then measured using an Instron 5585H universal testing machine (Instron Corporation), and values were calculated from the stress– strain curve within the 20–50% compressive strain region (**Extended Data Fig. 2d**).

**Extended Data Fig. 1.**
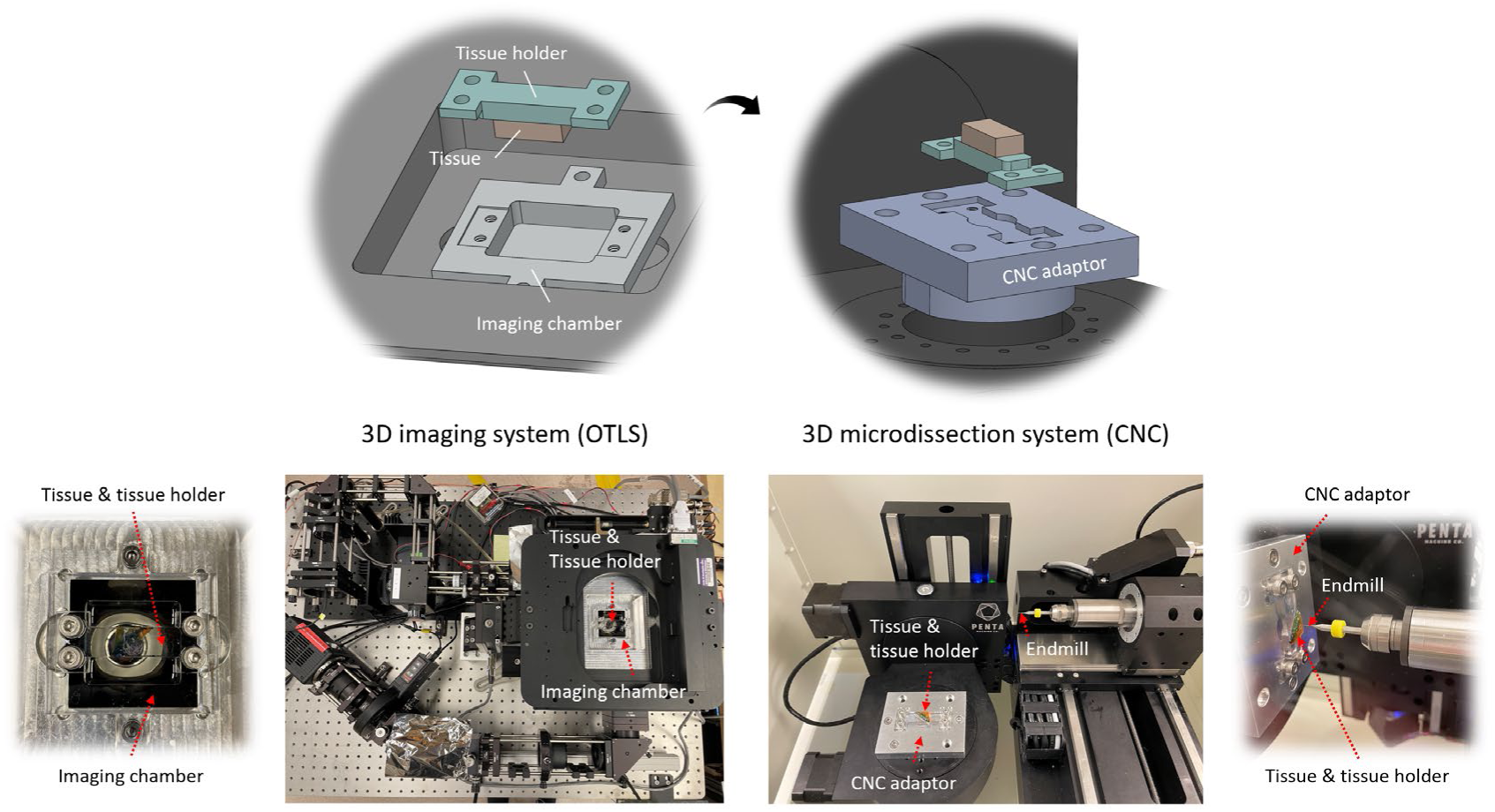
Schematics and photos of the open-top light sheet (OTLS) microscopy and 5-axis CNC-based microdissection system. Upper panel: CAD models of the imaging chamber in the OTLS microscope and the CNC adaptor on the 3D microdissection system, with the tissue holder fitted into matching slots in both. Bottom panel: photographs of the OTLS system and the portable 5-axis CNC machine, including a close-up of the imaging chamber and microdissection being performed on tissue.

**Extended Data Fig. 2.**
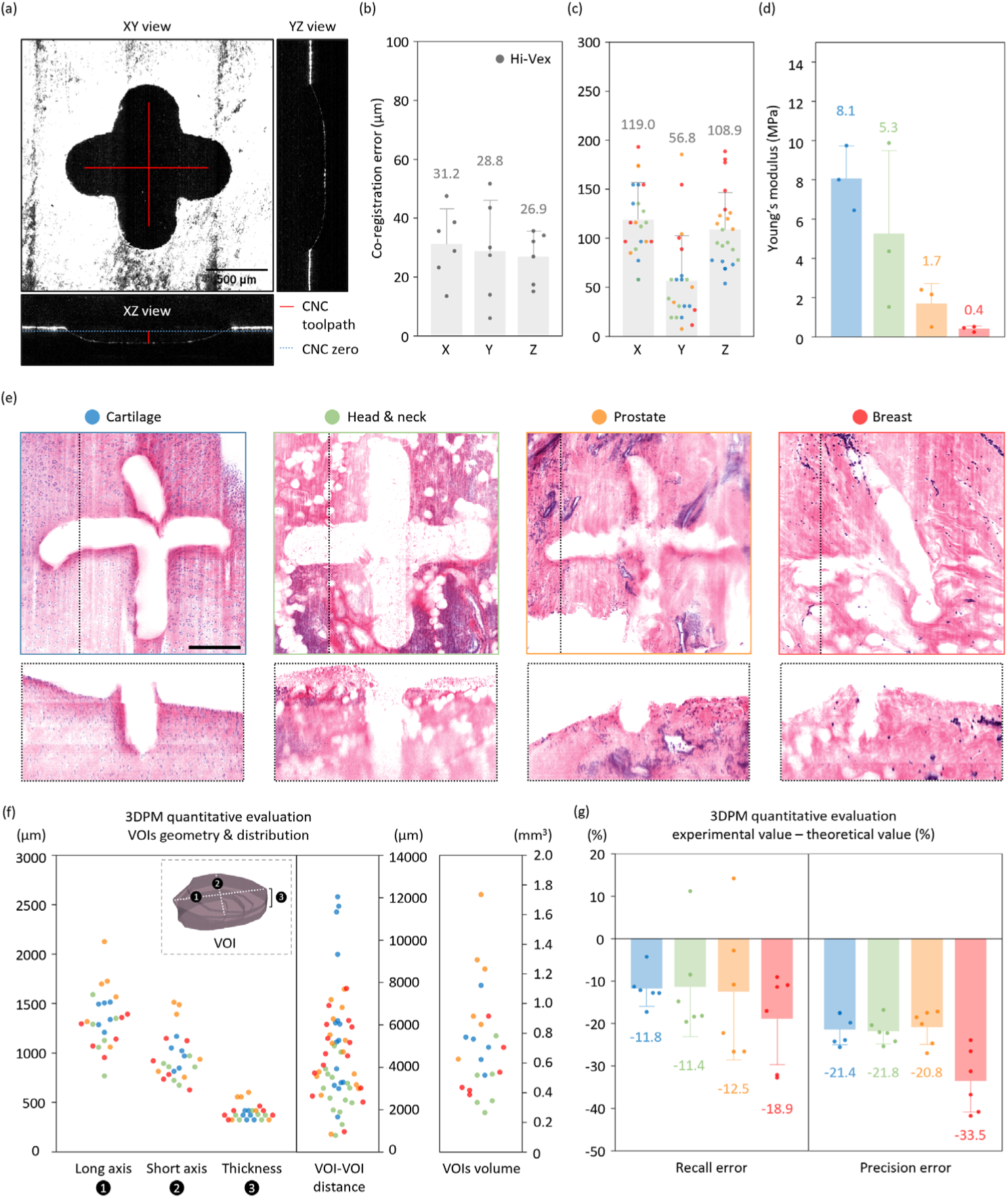
(a) Target cross-shaped pattern for co-registration between 5-axis CNC and OTLS systems. Note that manual CNC zeroing may be inaccurate; the actual CNC zero can be inferred by comparing the assigned depth with the actual cut depth. (b) Co-registration errors on Hi-Vex (rigid plastic) target materials, representing the highest achievable co-registration performance. (c) Co-registration errors for diverse tissue types: breast (red), prostate (orange), head and neck (green), and cartilage (blue). (gray bars: average across four tissue types). (d) Young’s modulus (stiffness) for the four tissue types. (e) Visualization of engraved cross shapes on different tissue types. The black line indicates the location of the cross-sectional views shown in the bottom row (scale bar: 300 µm) (f) Dimensions and distribution of VOIs used for the quantitative evaluation of 3DPM. The long and short axes were measured by projecting the 3D VOI onto the xy plane and fitting an ellipse. Thickness is the axial dimension of the VOI. VOI–VOI distance refers to the pairwise centroid-to-centroid distance (15 pairs per tissue type). (g) Differences in experimental and theoretical (toolpath-optimized) recall and precision metrics across the four tissue types.

**Extended Data Fig. 3.**
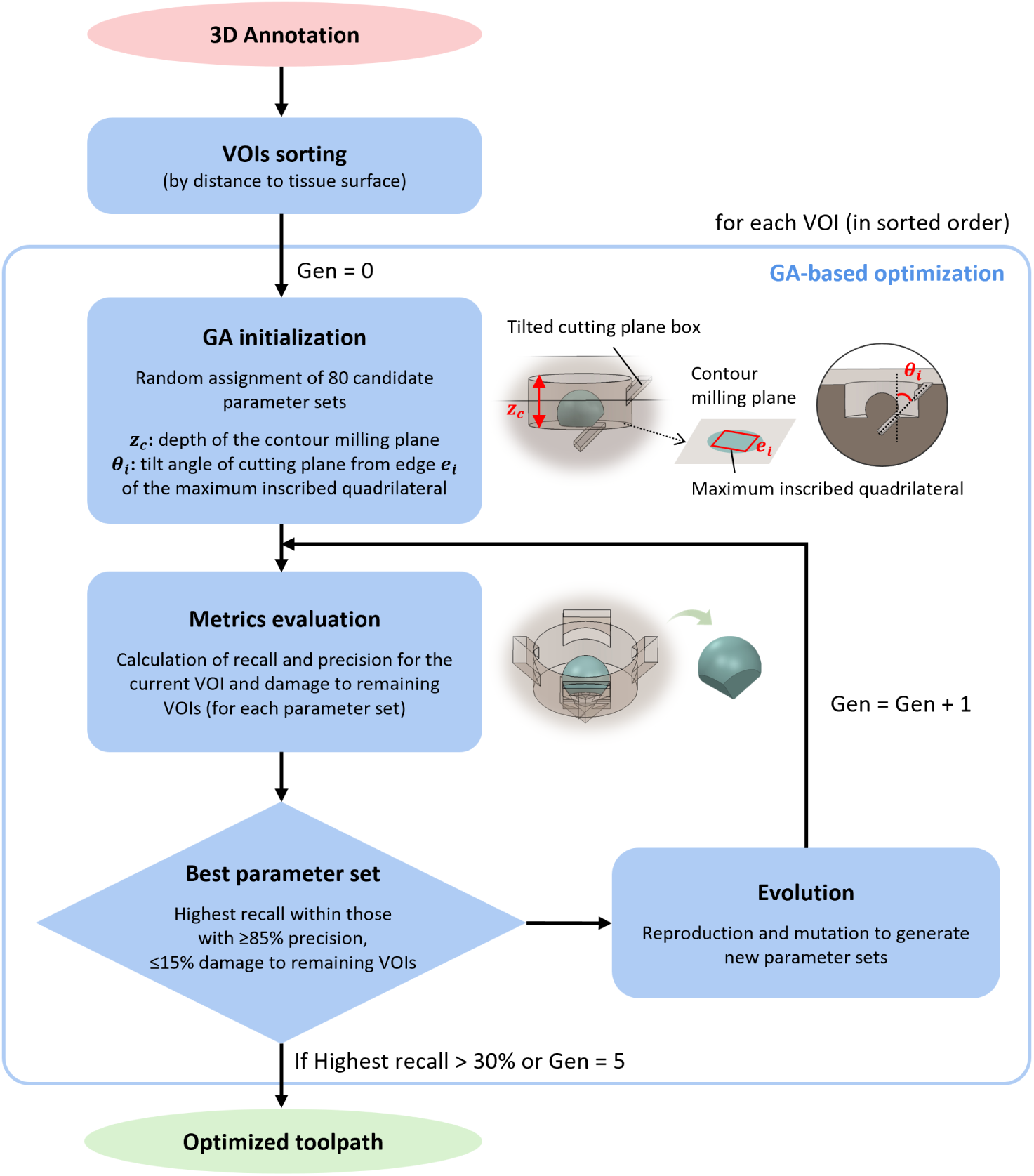
Schematic of the 3D microdissection toolpath optimization. VOIs are first sorted to determine the order of 3D microdissection. Each VOI then undergoes sequential optimization using a combination of random search and a genetic algorithm (GA).

**Extended Data Fig. 4.**
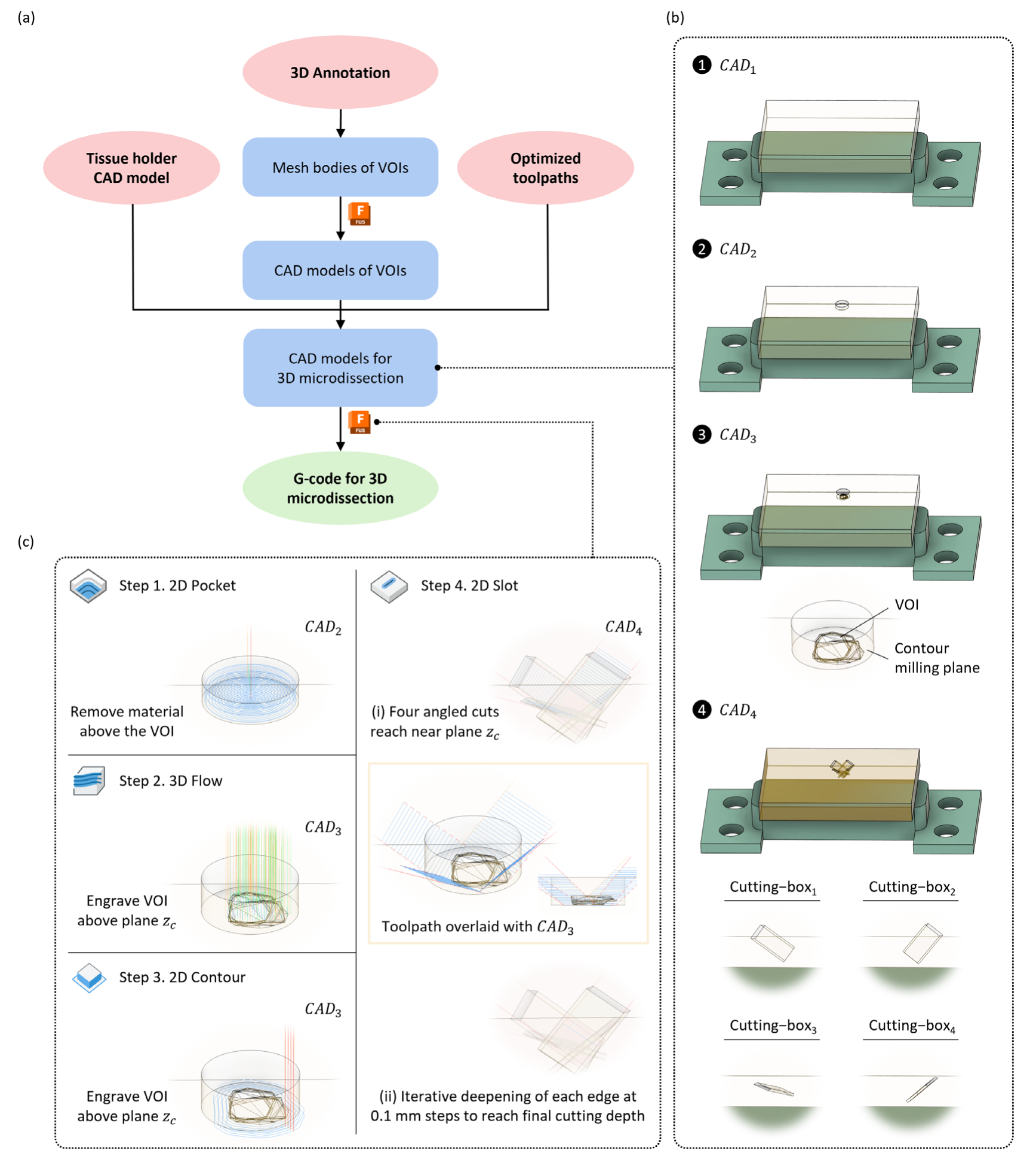
(a) Schematic of the 3D microdissection toolpath-generation workflow. The optimized toolpath results, together with the original 3D VOI annotations and the tissue-holder model (with preset positions), are input into a custom Python script to create CAD models for toolpath generation. Note that CAM software (Autodesk Fusion 360) is required to convert mesh bodies of VOIs into CAD-compatible formats. (b) Examples of four CAD models required for 3D microdissection toolpath generation. (c) Four steps of toolpath design and the corresponding algorithms used in Autodesk Fusion 360 (overlaid lines represent toolpaths, blue: cutting; red: ramp; green: lead-in/out; yellow: retract)

**Extended Data Fig. 5.**
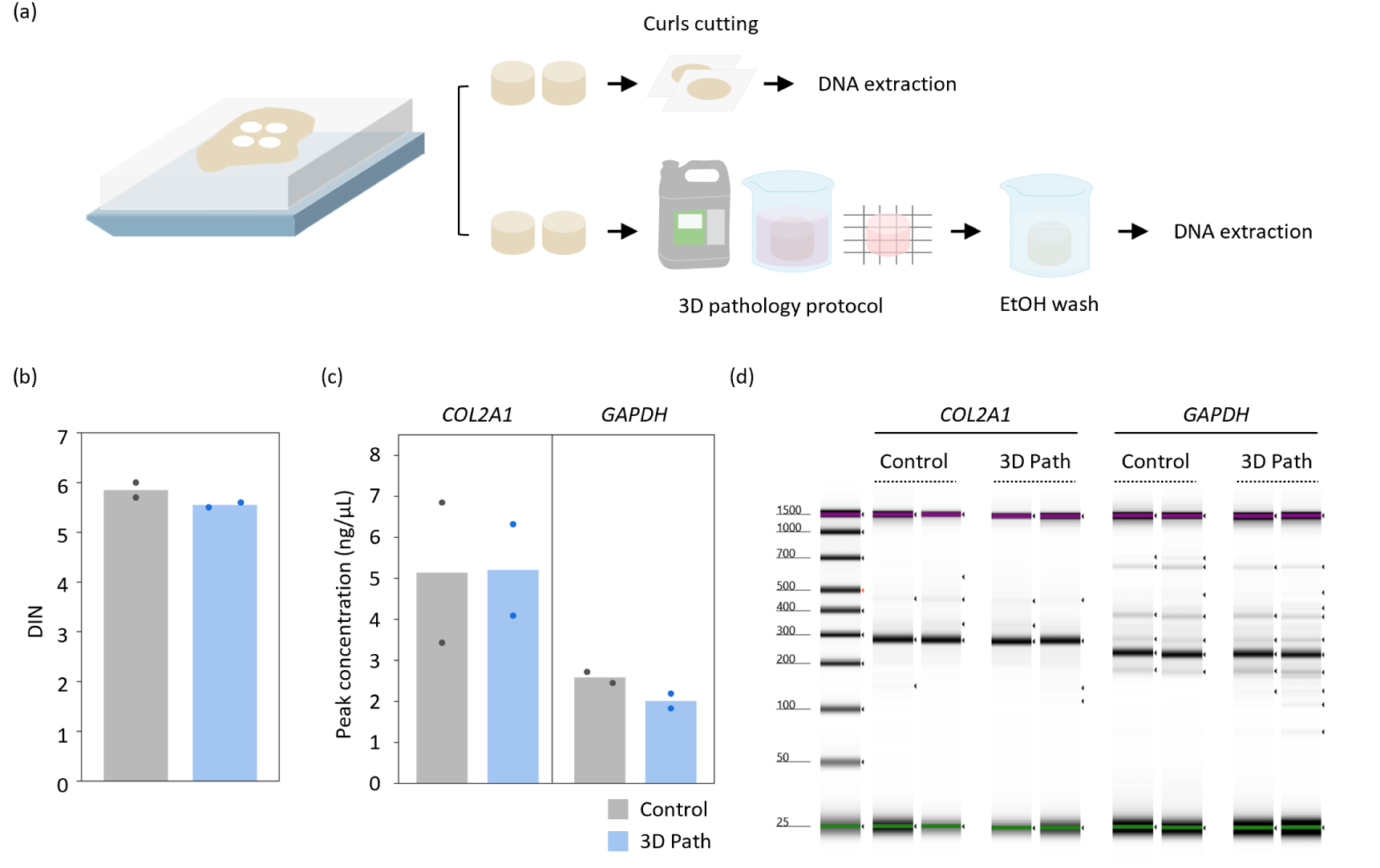
(a) Schematic of the validation workflow. Two punches underwent standard curl preparation and DNA extraction for FFPE tissues, while the other two were processed through the 3D pathology workflow followed by the modified DNA extraction protocol. (b) DIN scores of extracted DNA (c) PCR peak concentrations (measured by TapeStation) for amplified amplicons of specific regions within *COL2A1* and *GAPDH* in control and 3D pathology (3D path) groups. (d) Simulated gel images from TapeStation analysis, showing comparable PCR performance (amplifiability) between control and 3D pathology groups.

**Extended Data Fig. 6.**
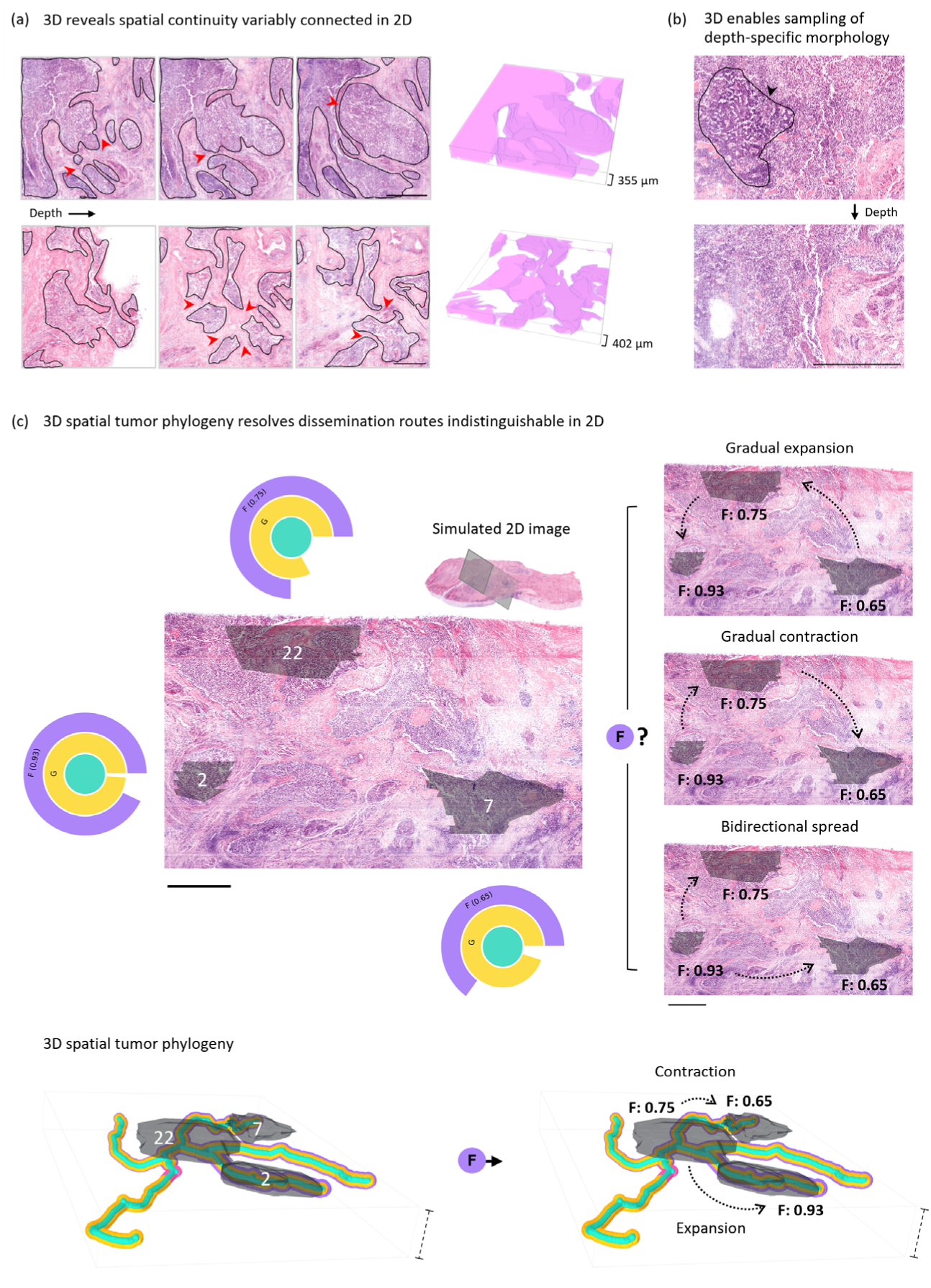
Importance of 3D information demonstrated in the prostate cancer phylogeny case: (a) The highly convoluted tumor architecture can only be fully characterized in 3D; the same foci, when viewed at different depths, may appear connected at some levels but disconnected at others (red arrows: regions seem to be disconnected in 2D). (b) Increasing the sampling ratio enables the capture of “rare” features — here, a trabecular growth region (black arrow) that is present only within a specific depth range. (c) Most importantly, 3D spatial information can elucidate possible subclone dissemination routes. As a contrasting example, a simulated 2D pathology image is provided, showing three VOIs that appear to be spatially disconnected. Without 3D spatial context, the separated tumor foci and the cancer cell fraction (CCF) of subclone F could support multiple possible dissemination routes (three shown here), each with different biological implications. (scale bar: 1 mm). In contrast, with 3D spatial tumor phylogeny, integration of tumor structure and CCFs across nodes can help to establish the most likely dissemination routes (scale bar: 1.2 mm).

**Extended Data Fig. 7.**
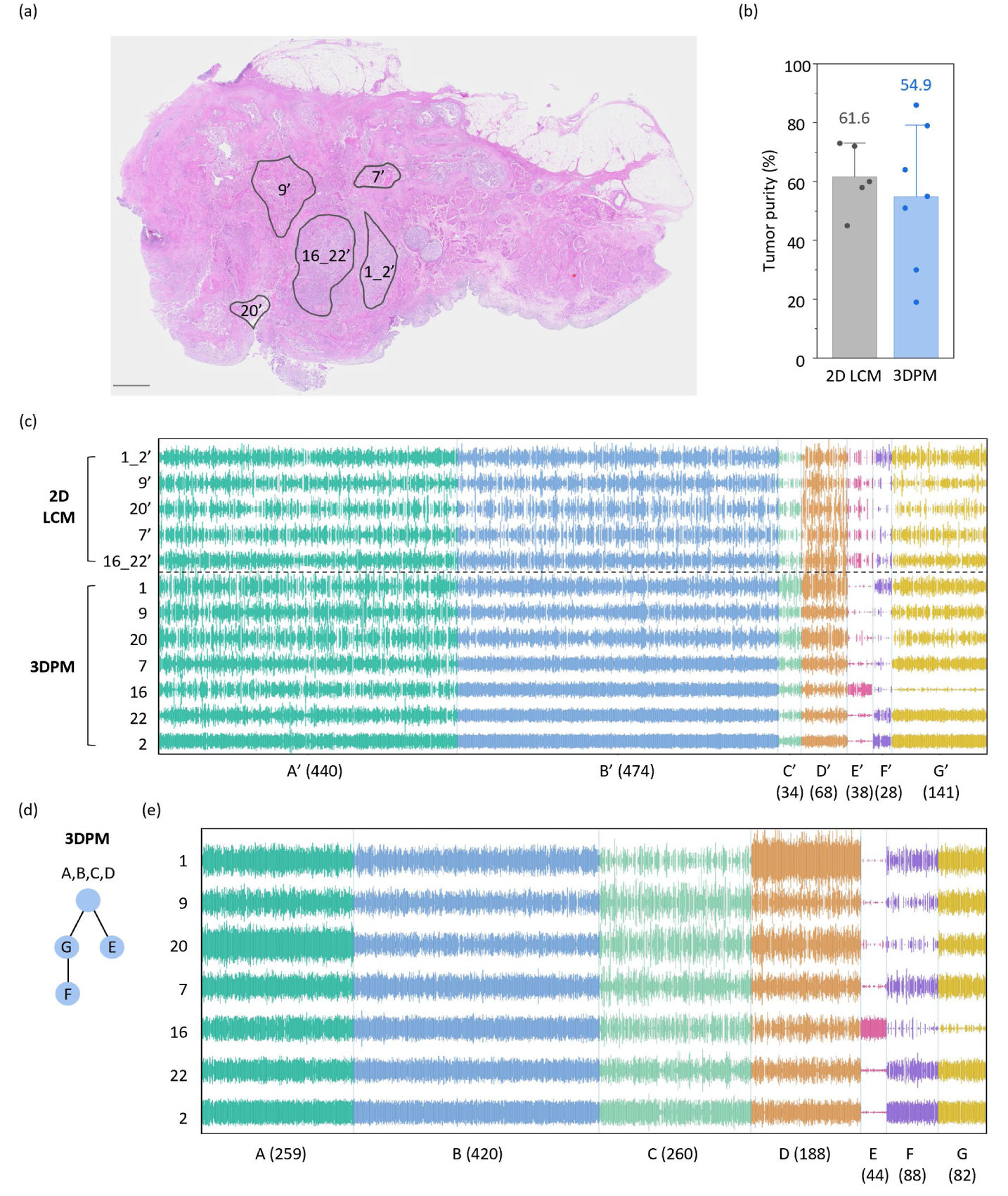
(a) Five tumor regions, corresponding to the locations of the 3D VOIs projected onto 2D slides cut from the top of the tissue block, were microdissected using 2D LCM. Since tumor morphology varies across sections, the regions on the 2D LCM slide were manually adjusted according to the actual morphology to ensure high tumor purity, and the final annotations were overlaid on the H&E image of an adjacent slide (scale bar: 2 mm). (b) Comparison of tumor purity between five regions (2D LCM) and seven regions (3DPM). No statistically significant difference was observed. (c) CCF-cluster plot from the joint analysis of 2D LCM and 3DPM samples. Each vertical line represents a SNV, with height corresponding to CCF; SNVs are grouped and colored by cluster. This comparison between methods demonstrated good agreement between the two approaches. (d) Phylogenetic tree and (e) CCF-cluster plot from the 3DPM-only analysis, corresponding to the phylogenetic tree shown in Figure 2. Here, the filtering criteria were optimized to retain high-confidence SNVs used for 3DPM-only analysis.

## Acknowledgements

This work was supported by funding from the National Institutes of Health (NIH) through R01EB031002 (J.T.C.L.), R01CA268207 (J.T.C.L.), R01DK138948 (J.T.C.L., W.M.G.), U54DK137328 (J.T.C.L.) and the Pacific Northwest Prostate Cancer SPORE P50CA97186 (L.D.T.). NIH support for W.M.G. includes U01CA152756, R01CA220004,637 U2CCA271902, U54CA163060, and U01CA182940. Funding for W.M.G. is also provided by the Prevent Cancer Foundation, Cottrell Family Fund, Evergreen Fund, and Listwin Foundation. Support was also provided by the Prostate Cancer United Kingdom (PCUK) charity through MA-ETNA19-005 (F.C.H., I.G.M, S.F., S.R.R., D.J.W., J.T.C.L.). Additional support was provided by the Department of Defense (DoD) Prostate Cancer Research Program: W81XWH-20-1-0851 (J.T.C.L.), W81XWH-18-10358 (J.T.C.L., L.D.T.), and the Advanced Research Projects Agency for Health (ARPA-H) D24AC00357 (J.T.C.L.). Support for K.W.B. is provided by National Science Foundation (NSF) Graduate Research Fellowship DGE-1762114. Any opinions, findings, and conclusions or recommendations expressed here are those of the authors and do not necessarily reflect the views of the NSF.

